# Worldwide occurrence records reflect a global decline in bee species richness

**DOI:** 10.1101/869784

**Authors:** Eduardo E. Zattara, Marcelo A. Aizen

**Affiliations:** Grupo de Ecología de la Polinización, INIBIOMA, Universidad Nacional del Comahue-CONICET, Quintral 1250, Bariloche (8400), Argentina

## Abstract

Wild and managed bees are key pollinators, providing ecosystem services to a large fraction of the world’s flowering plants, including ∼85% of all cultivated crops. Recent reports of wild bee decline and its potential consequences are thus worrisome. However, evidence is mostly based on local or regional studies; global status of bee decline has not been assessed yet. To fill this gap, we analyzed publicly available worldwide occurrence records from the Global Biodiversity Information Facility spanning more than a century of specimen collection. We found that after the 1980’s the number of collected bee species declines steeply, and approximately 25% fewer species were reported between 2006 and 2015 relative to the number of species counted before the 1990’s. These trends are alarming and encourage swift action to avoid further decline of these key pollinators.

## Introduction

Insects are the most specious group of animals and are estimated to encompass a large fraction of the Earth’s living biomass^1^. Given their historical abundance and ubiquity, along with the many familiar examples of extreme resilience to natural or intentional extermination, some insects have been traditionally viewed as the ultimate survivors of most apocalyptic scenarios. However, in the last two decades, a series of high-profile reports based mostly on local or regional evidence have repeatedly warned of a significant decline in insect diversity and biomass and raised the alarm about the potential consequence of this decline for the delivery of many ecosystem services ^2–5^. Among affected ecosystem services is plant pollination: insects are the main vectors for pollen transfer of most wild and crop flowering plant species ^6–10^. Bees (Hymenoptera: Apoidea: Anthophila), a lineage that includes about 20,000 described species, are the most important group of insect pollinators ^11,12^. Wild bee species are not only key to sexual reproduction of hundreds of thousands of wild plant species ^7^, but also to the yield of about 85% of all cultivated crops ^6,13,10^. There is mounting evidence that a decline in wild bee populations might follow or even be more pronounced than overall trends of insect decline ^12,14–17^. Such differential vulnerability might result from a high dependence of bees on flowers for food and a diversity of substrates for nesting, resources that are greatly affected by land conversion to large-scale agriculture, massive urbanization, and other intensive land uses ^18–20^. However, most studies on “bee decline” to date are based on local-, regional- or country-level datasets, and have a strong bias towards the Northern Hemisphere, particularly North America and Europe, where most long-term research projects capable of generating multidecadal datasets have been conducted ^4,12,21,22^.

To find an alternative approach to assess whether bee decline is a global phenomenon, we resorted to the data publicly available at the Global Biodiversity Information Facility (GBIF)^23^. The GBIF collects and provides “data about all types of life on Earth” from “sources including everything from museum specimens collected in the 18th and 19th century to geotagged smartphone photos shared by amateur naturalists in recent days and weeks”^23^. GBIF ingests data from a widely diverse range of data sources, localities, recording strategies, geographic areas, sampling intensities, etc., with each data source potentially plagued by both systematic and idiosyncratic biases ^24–27^. Although usage of GBIF data has been strongly criticized due to its inherent biases ^21,24,28–30^, most criticisms are usually aimed towards using its occurrence data to reconstruct and model species’ distribution ranges. Reconstructing geographic ranges and abundances from such “messy” datasets is indeed challenging. However, a binning approach in which a simpler question (“has a species been recorded anywhere during a given period?”) yields a yes/no answer can potentially be much more robust to sampling effort heterogeneity and geographic uncertainty ^31^. We reasoned that if bees are experiencing a global decline in the last few decades, then a generalized decrease in population size and range would result in increased rarity, diminished chance of observation and collection and, consequently, a diminished number of total species being observed and recorded worldwide each year.

## Results and Discussion

To test our hypothesis of global bee decline, we queried GBIF for all occurrence records of Hymenoptera prior to 2020 with either “Preserved specimen” or “Human observation” bases of record ^32^ (see Methods section below). Records of preserved specimens originate in vouchered collections such as those from museums and universities, or associated with biodiversity surveys and molecular barcoding initiatives, among others. Human observations, on the other hand, are records in which a given species was observed, but no voucher was collected; this category of records has been growing exponentially since citizen science initiatives became increasingly popular^33^. Because the preserved specimen records are likely to represent the most taxonomically traceable source of information within the GBIF dataset^33,34^, we made parallel analyses for both the full dataset and the specimens-only subset. We filtered the datasets to six families of the superfamily Apoidea that conform the Anthophila or “true bees”: Melittidae, Andrenidae, Halictidae, Colletidae, Megachilidae and Apidae (we excluded the small family Stenotritidae from our analysis, since it has only about 21 species restricted to Australia) ^11^.

Plotting the total number of records per year in both datasets show that the number of worldwide bee occurrence records follows a mostly monotonic increasing trend that becomes steeper after 1990 (Fig. 1A). Since the four most recent years (2016-2019, marked with * in Fig. 1A) show a noticeable drop in records, likely due to time lags in data entry ^35^, we excluded those years from further analyses to avoid a downward bias in most recent years. In contrast, while the number of recorded species per year during the same period also increases initially, it reaches a steady maximum after 1950 but then shows a noticeable decline starting near the end of the 20th century (Fig. 1B). This negative temporal trend persisted even when number of records and of contributing collections, institutions and datasets are considered (generalized least squares estimate ± s.e. for the period 1986-2016: −31.9±11.0, *t* value: −2.9, *p* = 0.008). Thus, fewer species have been reported globally within GBIF records since approximately the 1990s.

**Fig. 1.**
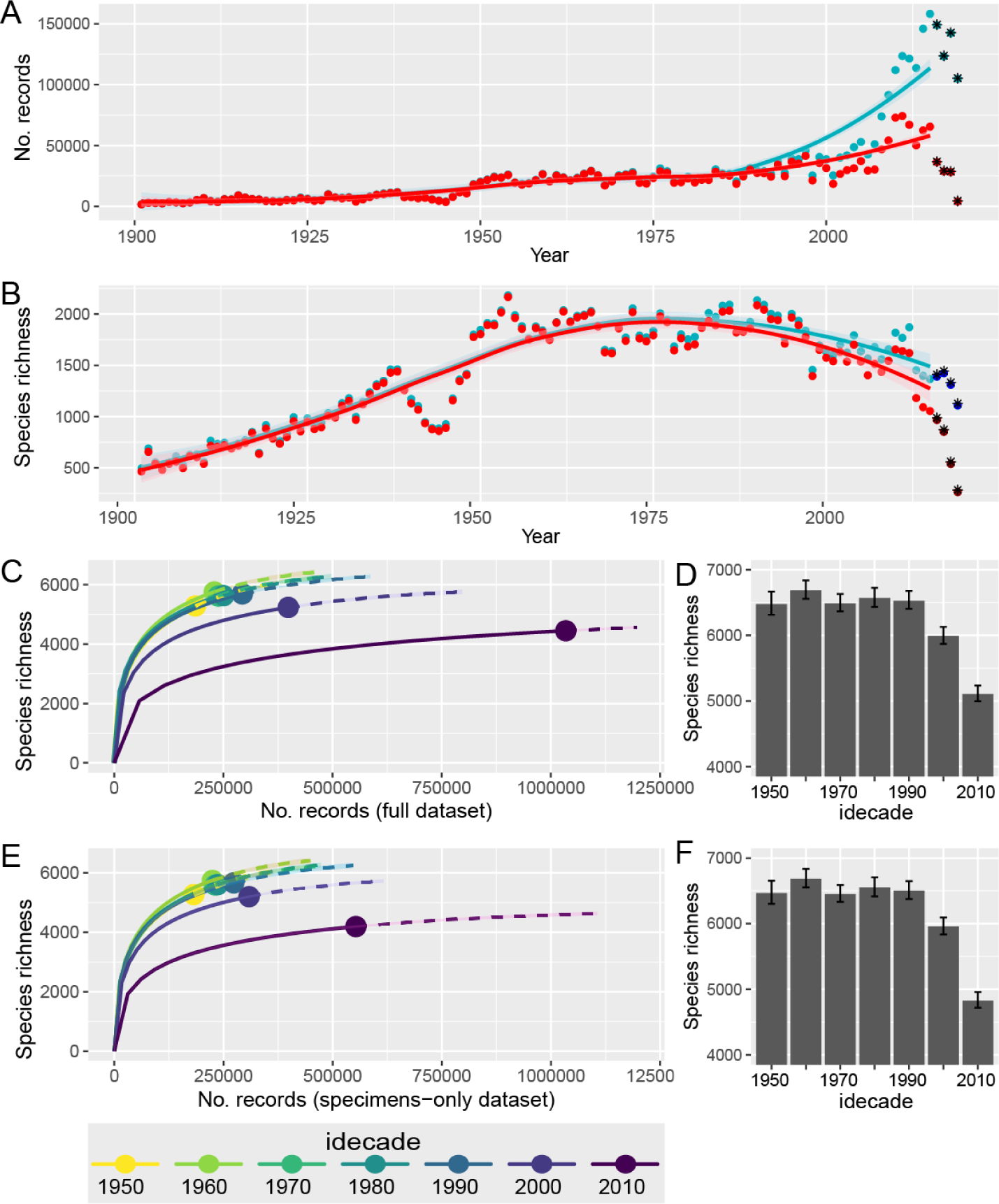
Despite increasing number of specimen records, the number of worldwide recorded bee species is sharply decreasing. (A) Number of worldwide GBIF records of Anthophila (bees) occurrences per year in the full (blue) and specimens-only (red) datasets. The curves represent loess fits with a smoothing parameter α = 0.75 up to 2015. The four most recent years (2016-2019, labeled with *) were excluded from further analysis. (B) Number of bee species found each year in the full (blue) and specimens-only (red) datasets. (C) Chao’s interpolation/extrapolation (iNEXT) curves based on the full dataset. Data were binned into ten-year periods (*idecades*) from 1946 to 2015. The symbols show actual number of specimen records and separate interpolated (left, full line) from extrapolated (right, dashed line) regions of each curve. (D) Values of the asymptotic richness estimator by idecade (see main text) for the full dataset (error bars mark upper and lower 95% confidence intervals). (E) Chao’s interpolation/extrapolation (iNEXT) curves based on the specimens-only dataset. (F) Values of the asymptotic richness estimator by idecade for the specimens-only dataset.

To remove potential biases introduced by year-to-year heterogeneity of data sources, we binned records every 10 years starting from 1946 (after the end of World War II, which caused a noticeable dip in collection intensity, see Fig. 1A) and until 2015 inclusive; we call these bins “idecades” and name them by the multiple-of-ten year in the middle. We then used rarefaction based interpolation/extrapolation curves (iNEXT) and asymptotic richness estimators ^36,37^ to compare idecadal changes in richness of species records. In this analysis, accumulation curves are very similar from the 1950’s to the 1990’s but flatten considerably to reach lower asymptotes for the 2000’s and 2010’s (Fig. 1C,E), again showing that the number of species among bee specimens collected worldwide is showing a sharp decline. More specifically, we found a reduction of about 8% during the 2000s in both datasets, and of 22% and 26% during the 2010s for the full and specimen-only datasets, respectively (Fig. 1D, F).

Bee families in our dataset are heterogeneous in term of richness and abundance, and the observed trends might be driven by just a few bee clades. To make a more phylogenetically-explicit analysis exploring whether bees show a differential temporal trend compared to their closest relatives, and whether particular bee families are more endangered than others, we re-analyzed the specimen dataset, this time retaining also records for two families of carnivorous apoid wasps, Crabronidae and Sphecidae, that are sister to Anthophila, and for another highly diverse, non-apoid hymenopteran family, the Formicidae (ants) ^38^. The results show different patterns of species richness in records of each family, with noticeable phylogenetic structure (Fig. 2). Long-tongued bees (Megachilidae and Apidae) show a steepening decline starting at 2000’s, while short-tongued bees show declines starting earlier (Andrenidae and Halictidae) or later (Colletidae). These declines in richness of recorded species relative to the average number found between 1950 and 1990 ranged from 17% for Halictidae to over 41% for Melittidae. Comparisons between Antophila families and two families of apoid wasps sister to bees, and to a more distantly related family, the true ants (Formicidae) revealed contrasting trends (Fig. 2). While both wasp families also show declining trends, they present different patterns than bees. Record richness of sphecid and crabronid wasps both show a smoother decrease initiating earlier than the 2000’s. In contrast, ants show very little evidence of global record richness decline, but rather a trend towards an increase in the number of recorded species. Although the limited number of bee families precludes a formal analysis of phylogenetic patterning, closely related families (e.g., Apidae and Megachilidae, or Colletidae and Halictidae) seem to share more similar trends in terms of timing and magnitude of species richness decline than less related families. This hint of phylogenetic patterning becomes even more apparent when considering the two apoid wasp families, Crabronidae and Sphecidae (Fig. 2). Interestingly, a very similar pattern – in which bees show a strong, recent decline, wasps show a gentler decline starting earlier, while ants remain steady – was recently reported using a quite different analytical approach on a substantially different and more geographically limited dataset ^39^. Altogether, family-specific trends and asymptotic richness estimates show that the overall decline in global bee record richness is not driven by any particular family. Instead, a generalized decline seems to be a pervasive feature within the bee lineage.

**Fig. 2.**
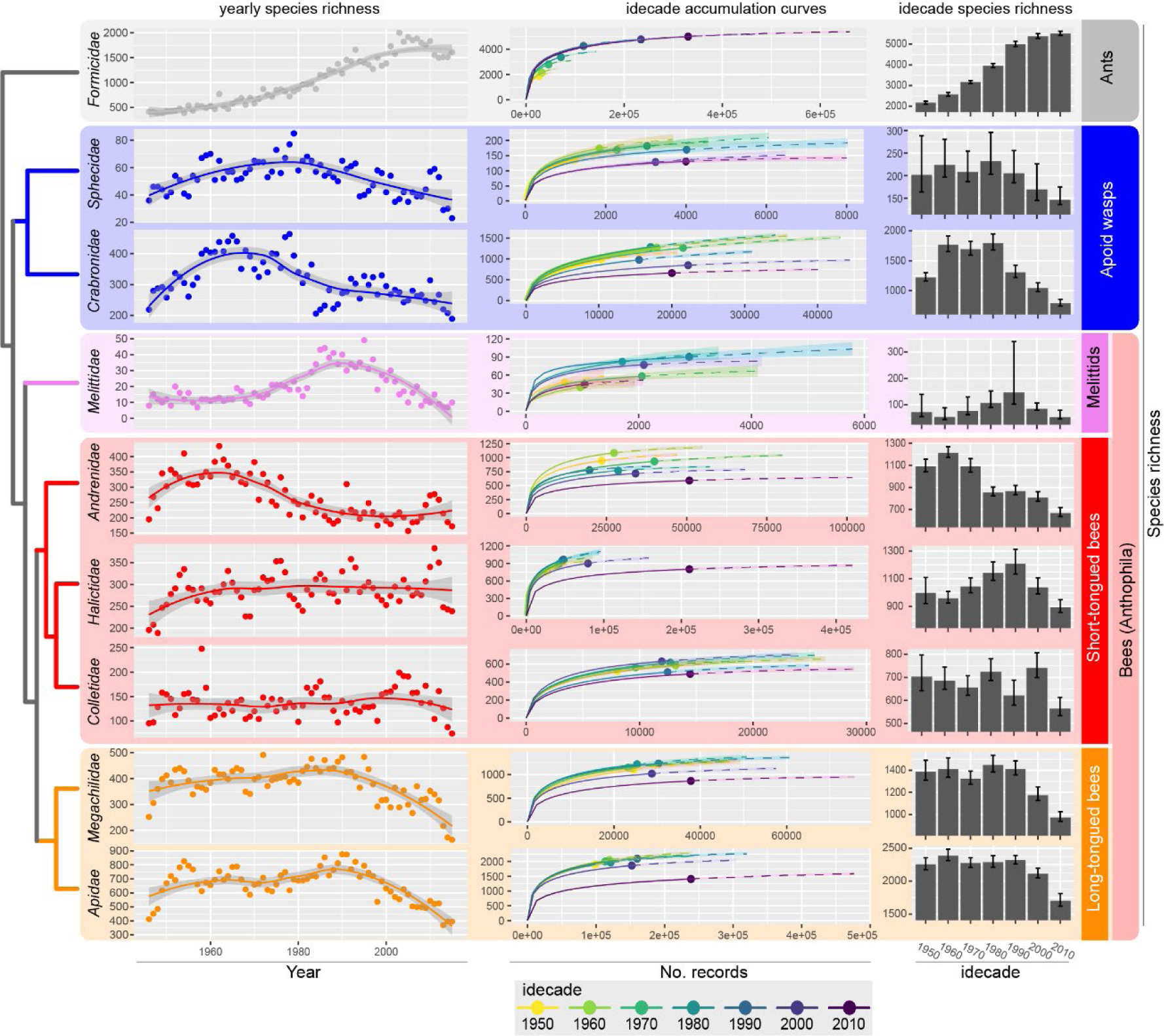
Decline patterns in worldwide records of bees are generalized but phylogenetically structured. Phylogenetic relationships among each of the six families of bees (Anthophila, lower six rows), two related families of non-flower associated apoid wasps (2nd and 3rd rows), and the less related, highly specious ant family (top row). The left row shows number of species per year in GBIF records from 1946 to 2015 based on the preserved specimen dataset – the curves represent loess fits with a smoothing parameter α = 0.75; the middle row shows Chao’s interpolation/extrapolation curves based on GBIF records, grouped by idecade for the period 1946-2015; the right row show the asymptotic estimates of richness by idecade for the same period (error bars mark upper and lower 95% confidence intervals).

To rule out the possibility that the method we used to estimate richness does not correlate with actual bee diversity, we compared the asymptotic estimator of total richness for each family based on GBIF records with the total known number of species and found a linear correlation between both estimates across families (Fig. S1). Another potential artifact causing a decline in recorded bee diversity in the last two idecades could be an increasing loss in taxonomic expertise during that period ^40–42^. Under such scenario, we would expect the fraction of records unidentified to the species level – a reasonable proxy for lack of expertise ^33^ – should have stayed approximately constant until the last two decades and then increased noticeably. While the fraction of records missing species’ identification shows an overall increase in the last 120 years, this trend has actually reversed since the 2000’s (Fig. S2). This result is consistent with previous analyses of the GBIF dataset^33^, and shows that potential loss of taxonomic expertise cannot explain the decline in bee record diversity seen at the last two decades.

Next, we explored the geographic distribution of the dataset, by sub-setting the data by continent and repeating the analyses. Overall, GBIF has a strong bias towards North American and European records ^35^, and this bias results in a very uneven contribution of each continent to decadal number of records (Fig. S3). North America (including Central America and the Caribbean) has the largest and most even representation of records across decades (between 46 and 75% of global records) and shows its steepest decline in species richness between the 1990’s and the 2010’s (Fig. S4). In contrast, Europe shows two separate periods of decline, one between the 1960’s and the 1970’s and a more recent drop between the 1980’s and 1990’s but stabilizes afterwards (Fig. S4). Africa shows a sustained fall in species richness since the 1980’s, whereas in Asia the decline seems to have started two or three decades earlier (Fig. S4). The trend in South America is less clear, although estimated richness also decreases in the last ten years of the dataset (Fig. S4). Overall, analyses of the dataset at a continental scale show heterogeneity in both the proportional and absolute contributions to the records, and in the timing and magnitude of the decline in species richness. However, despite large differences in data availability and, perhaps, except for Oceania, a decline in species richness of bee records seems to be common to all continents.

A global decline in bee record diversity could relate to a proportional decrease in bee abundance, so that rare species become rarer or even extinct, and abundant species become less abundant. Alternatively, the less abundant species could be declining strongly, whereas abundant species might be declining at a lower rate or even thriving. These different scenarios are expected to leave a distinctive signature in the temporal pattern of relative record abundances. Under the first scenario, the sharp decrease in species richness estimates should not be accompanied by a decrease in evenness, a measure of how equally total record abundance is partitioned among species, whereas under the second scenario there should be a parallel decrease in record evenness. As expected from the hypothesis of an abundance-related differential species decline, plotting Pielou’s index (a common measure of evenness ^43^) per year of bee records shows a strong decreasing trend since the 1990’s for both datasets (Fig. 3). Therefore, this decline in species richness of records can relate to either a global change in how an invariant bee diversity is sampled, leading to more infrequent reporting of many species and much more frequent reporting of a few other species, or to a global phenomenon by which thousands of species are becoming too rare to be sampled while fewer species are becoming dominant and perhaps even increasing in abundance. These two alternatives are not mutually exclusive, and both increased sampling and reporting bias and declining bee biodiversity should be a matter of concern.

**Fig. 3.**
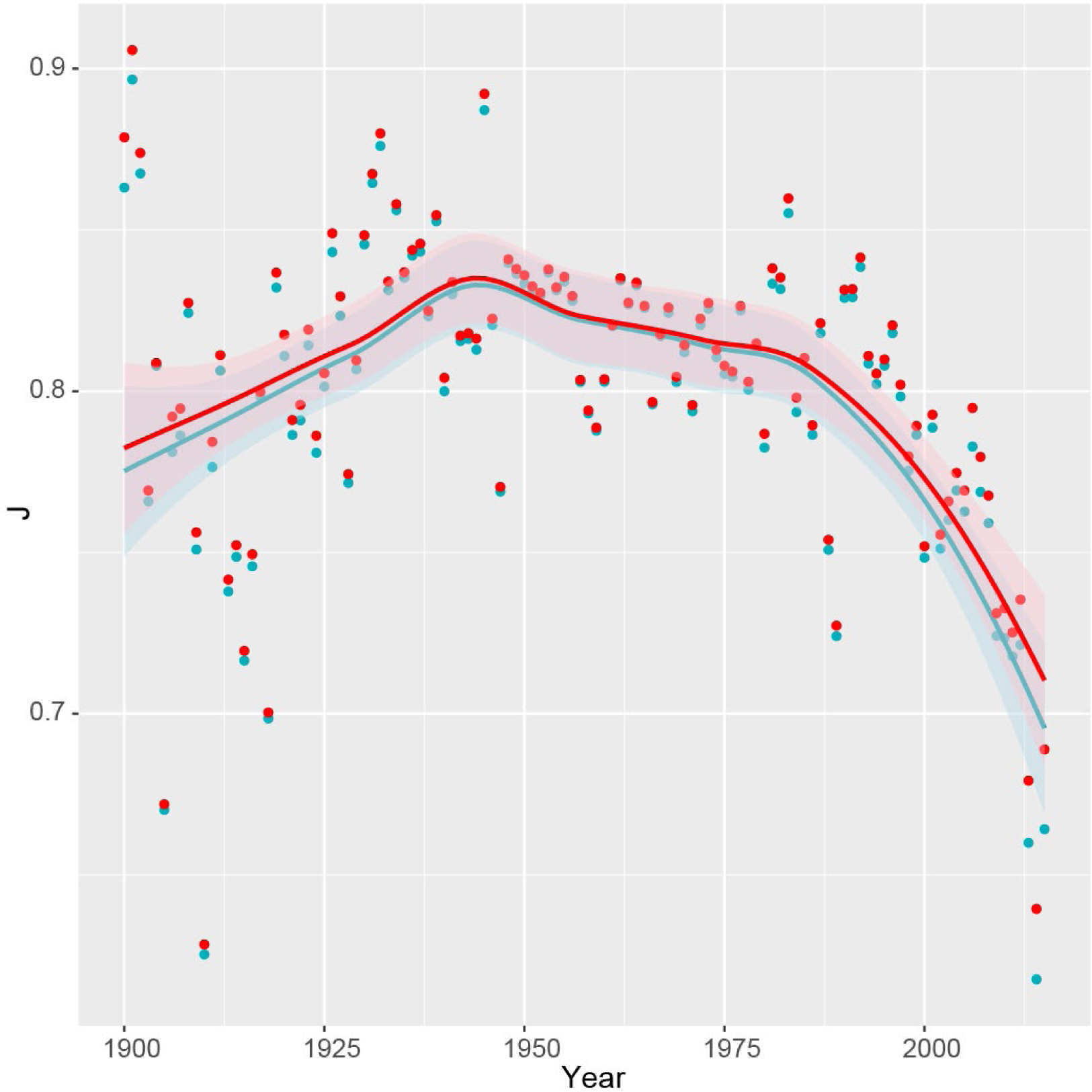
Overall representation of worldwide bee species based on global records is becoming increasingly uneven over time. Estimate of Pielou’s index of sample evenness per year in the full (blue) and specimens-only (red) datasets from 1900 to 2016. The lines show respective loess fit curves with a smoothing parameter α = 0.75.

Our results support a hypothesis of overall decline in bee diversity at a global scale. If trends in species richness of GBIF records are reflecting an actual trend in bee diversity, then this decline seems to be occurring with distinctive characteristics in every bee family and in most continents. Interestingly, this trend appears to be a relatively recent phenomenon that accentuated in the nineties, at the beginning of the globalization era, and continues to the present. The globalization era has not only been a period of major economic, political and social change, but also of accelerated land-use transformation ^44^. Bees thrive in heterogeneous habitats, even those driven by man ^18,45^, where they find a diversity of floral and nesting resources. However, land devoted to agriculture, particularly to monoculture, has expanded in several regions of the world since the 1990s ^44^. This has led not only to higher habitat homogeneity, which can relate by itself to more impoverished and spatially homogeneous bee assemblages ^18,46^, but also to higher use of pesticides and other agriculture chemical inputs that have direct and indirect lethal and sub-lethal effects on bee health ^47^. Effects of climate change on shrinking bee geographical ranges have been also documented in Europe and North America ^4^. Lastly, a booming international bee trade has involved the co-introduction of bee pathogens, that may cause bee decline, like the emblematic case of the giant Patagonian bumble bee, *Bombus dahlbomii* ^48^. These drivers can act synergistically, which can have accelerated a process of bee decline. Phylogenetic patterning in the trend of recorded species diversity among the different bee families (Fig. 2) suggests that different lineages can be differentially affected by different drivers, likely based both on their common geographical distribution and shared clade-specific biological and ecological traits ^21,49,50^.

Associated with the declining trend of richness of species records is a trend of increasing dominance of records by a few species. Increasing dominance by one or a few species can be observed at the regional scale, like the case of invasive *Bombus terrestris* in southern South America ^51^ or the western honeybee *Apis mellifera* in the Mediterranean ^52^. The western honeybee has been introduced in every single continent from its original geographical range in Europe and Africa. Although both domesticated and wild populations of the western honeybee seem to be declining in several countries, this species is still thriving globally ^53^. Correspondingly, an increasing fraction of the total global bee records is composed by *Apis mellifera* occurrences (Fig. S5). A consequence of increasingly less diverse and uneven bee assemblages could be an increase in pollination deficits, causing a reduction in the quantity and quality of the fruits and seeds produced by both wild and cultivated plants. Less diverse bee assemblages at both local and regional scales have been associated with lower and less stable yields of most pollinator-dependent crops ^13^.

GBIF is certainly not a source of systematically collected data, and this should be borne in mind when interpreting the results of our analyses ^21,27,35,54,55^. Spatial and temporal biases in collection intensity (e.g., targeted programs might enrich the abundance of specific species/groups at specific spans and regions) can generate spurious trends. In our analysis, we counted every species only once per year regardless of how many records it had for a given year; this filters out biases due to sporadic intensive sampling campaigns. Biases introduced due to targeted collection efforts or local/regional events (e.g., changes in research and conservation policies, economic downturns, social unrest, etc.) are likely, yet most such biases tend to be spatially and temporally restricted, and are less likely to systematically affect trends at the global, multi-decadal scale of this analysis. Indeed, several potential biases would be expected to deflate, rather than inflate our results. For example, collectors targeting rare species would be expected to enrich the number of species (unless many species are becoming so scarce that they just cannot be found).

Nonetheless, our continent-level analysis showed that regions with the best temporal and spatial coverage (i.e., Europe and North America, Fig. S3) are the ones showing the clearest signal for decline (Fig. S4); our results agree with several existing reports at local, national and subcontinental level ^14,16,17,39,56–61^. Furthermore, none of those biases can explain the noticeable phylogenetic contagion seen in the trends (Fig. 2) better than the fact that the hymenopteran groups we analyzed have a considerable phylogenetic signal in their ecology and life history traits and would be expected to show phylogenetic clustering in their response to drivers of decline ^50^.

Unsurprisingly, when data is disaggregated by country, agreement between country-level results and existing reports improves as the number of records increases. For example, our data reflects a clear and continuous decline in bee diversity in the USA ^56,57,60^ (with over 1 million records), a decline in Brazil^61^ during the last two decades (∼190k records), but shows no clear loss of richness in Great Britain (∼25k records), or much uncertainty in an apparent trend in bee species loss in Panama (∼9k records), despite reports of bee decline in all those countries^14,16,17,62^ (Fig. S6). Interestingly, reports on decline of British bees are based on occurrence data that is not publicly available – i.e. ∼300k records from the Bees, Wasps and Ants Recording Society (BWARS: http://www.bwars.com/). This suggests that, besides data source heterogeneity, a major source of bias and inaccuracy of results derived from GBIF data result from obstacles to data mobilization, and highlights the need to increase efforts to remove barriers to data sharing and discourage funding agencies from allowing data sequestration.

Thus, while the inherent heterogeneity and biases of aggregated datasets as those offered by GBIF make them unreliable as a direct (i.e., unfiltered, uncorrected) data source of predictive models, they can still be used within a hypothesis-driven framework to test whether bees (or any other taxon) as a group are declining worldwide. In this context, our results largely agree with the hypothesis that current regional reports of declining bee diversity reflect a global phenomenon.

## Conclusions

One of the most important pieces of missing information of the global report on Pollinators, Pollination and Food Production of IPBES ^63^ was the lack of data on global bee decline, despite the many local and few regional reports pointing out that this decline could add to a global phenomenon. Despite all its shortcomings, GBIF still is probably the best global data source available on long-term species occurrence and has the potential to contribute in filling this critical knowledge gap. Its analysis supports the hypothesis that we are undergoing a global decline in bee diversity that needs the immediate attention of governments and international institutions. Under the most optimistic interpretation – that bees are not declining, and the trends we find are an artifact of data collection - our results would indicate that global efforts to record and monitor bee biodiversity are decreasing over time. However, and given the current outlook of global biodiversity ^4,5,10,12^, it is more likely that these trends reflect existing scenarios of declining bee diversity. In the best scenario, this can indicate that thousands of bee species have become too rare; under the worst scenario, they may have already gone locally or globally extinct. In any case, a decline in bee diversity driven by either increasing rarity or irreversible extinction will affect the pollination of wild plants and crops, with broader ecological and economic consequences. Slowing down and even reversing habitat destruction and land-conversion to intensive uses, implementation of environmentally friendly schemes in agricultural and urban settings, and programs to re-flower our world are urgently required. Bees cannot wait.

## Methods

### Datasets

An initial query at the database of occurrence records at the Global Biodiversity Information Facility (www.gbif.org) using the filters [Scientific Name = “Hymenoptera”, Basis of Record = “PRESERVED_SPECIMEN” | “HUMAN_OBSERVATION”, Year <2020] resulted on 9,176,688 total records involving 2,374 datasets ^32^. Data were downloaded as a text file and filtered for records identified to species levels and belonging to Anthophila (defined as the families Melittidae, Andrenidae, Halictidae, Colletidae, Megachilidae and Apidae; 3,459,086 records). We also retrieved records for two closely related families of apoid wasps (Crabronidae and Sphecidae; 283,331 records), or the true ants (Formicidae; 1,121,857 records). Phylogenetic relations between all these nine families follow recent phylogenomic results ^38^.

### Analyses

All datasets were analyzed using a customized script written and executed within the R computing environment ^64^. The complete annotated script is available as Supplementary Materials, and can be used to fully reproduce all results, or adapted to re-run the analyses on other datasets. Data was processed using the tidyr^65^, dplyr^66^ and data.table^67^ packages.

After removing records without “year” data, yearly counts of records and species were plotted using ggplot2^68^. Significance of a negative trend was tested by fitting yearly counts of records, species, collections, institutions and datasets a generalized least squares model with the formula `sp ∼ year + records + collections + institutions + datasets`, with an autoregressive-moving average autocorrelation structure of order (1,0). Then, each year was assigned to a 10-year period termed “idecade” (for inter-decade), corresponding to a regular decade shifted four years into the past (e.g, the 1990’s idecade spans 1986 to 1995). Records by species and idecade were counted and stored in a matrix of *m* species × *7* idecades (1950’s to 2010’s). This matrix was used as abundance data input for the `iNEXT` function of the iNEXT package ^37^ to estimate rarefaction-based interpolation/extrapolation (iNEXT) curves and Chao1 asymptotic estimators of species richness ^36^. We also compared the asymptotic estimator for species richness for each family with the total number of species listed for each family in the taxonomic framework of the Integrated Taxonomic Information System (www.itis.gov).

To estimate potential biases caused by changes of taxonomic expertise over time, we re-filtered the initial GBIF query without excluding records without a species ID, then counted the number of records with or without a species id per year ^33^. To analyze trends at continental level, we added a “Continent” field to the base dataset via table joining to a list of countries, country codes and continents from https://datahub.io/JohnSnowLabs/country-and-continent-codes-list. We then repeated the analyses splitting the dataset by continent. Continent and country-specific shapes were taken from https://github.com/djaiss/mapsicon. To show trends in equitability of species abundance across records over time, we calculated Pielou’s evenness index ^43^, *J=*Σ*p*_*i*_ln(*p*_*i*_)*/*log(*S*) for *i=*1 to *S*, the total number of species, for each year between 1900 and 2018, using the diversity functions from the package `vegan`^69^. The contribution of a given species (e.g., *Apis mellifera*) was calculated as yearly number of the species records divided the total number of records for that year and plotted as a function of year.

## Acknowledgments

We thank Lawrence Harder, Lucas Garibaldi, Gherardo Bogo, Reto Schmucki, Ignasi Bartomeus, Castor Guisande and several other colleagues who provided both encouragement and constructive criticism to earlier drafts of this manuscript, and to the GBIF Secretariat for maintaining this great resource. This project was inspired by work done at the Safeguarding Pollination Services in a Changing World (SURPASS) workshops that took place at Puerto Blest, Bariloche, Río Negro, Argentina in 2018, and Seaton, Devonshire, UK in 2019, supported by the Researcher Links Workshop grant, ID 2017-RLWK9-359543120, under the UK LATAM partnership funded by the UK Department of Business, Energy and Industrial Strategy (BEIS) and Argentina’s CONICET, and delivered by the British Council.

## Footnotes

### Author contributions

Conceptualization, E.E.Z. and M.A.A.; Data Curation: E.E.Z.; Formal analysis: E.E.Z. and M.A.A.; Visualization: E.E.Z.; Writing – original draft, E.E.Z. and M.A.A., Writing – review & editing, E.E.Z. and M.A.A.

### Data and materials availability

Occurrence record data used in this paper can be downloaded from https://doi.org/10.15468/dl.ysjm4x; original sources traceable via GBIF.org. The R language script used to analyze the data and generate the plots is available at https://github.com/ezattara/global-bee-decline.

## Supplementary Figures

**Figure S1:**
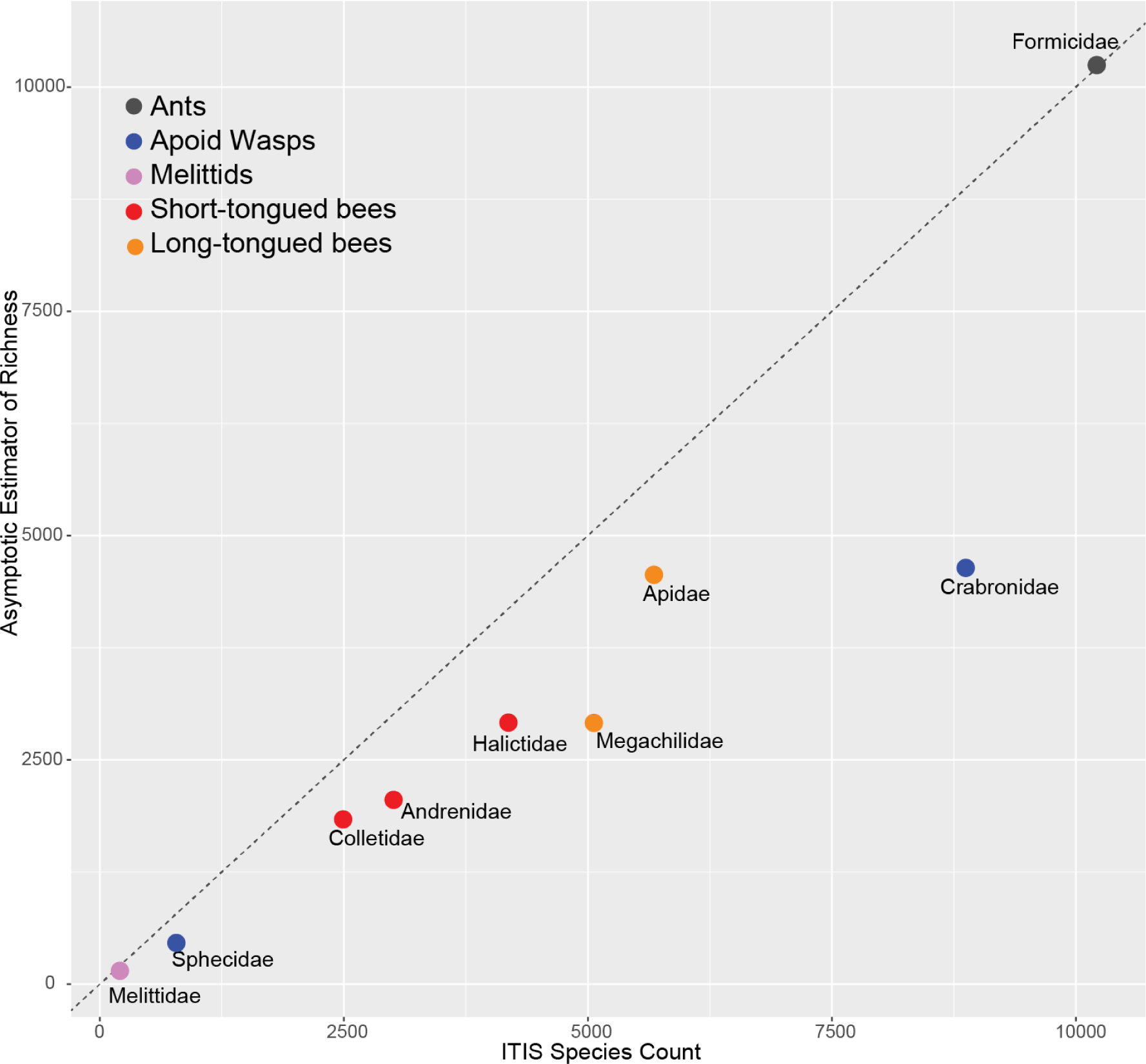
While often underestimating the known richness of each family, Chao’s asymptotic estimators of species richness based on all-times GBIF global records of preserved specimens show a linear correlation with actual species diversity. The dotted line shows the identity diagonal. ITIS stands for Integrated Taxonomic Information System (www.itis.gov).

**Figure S2:**
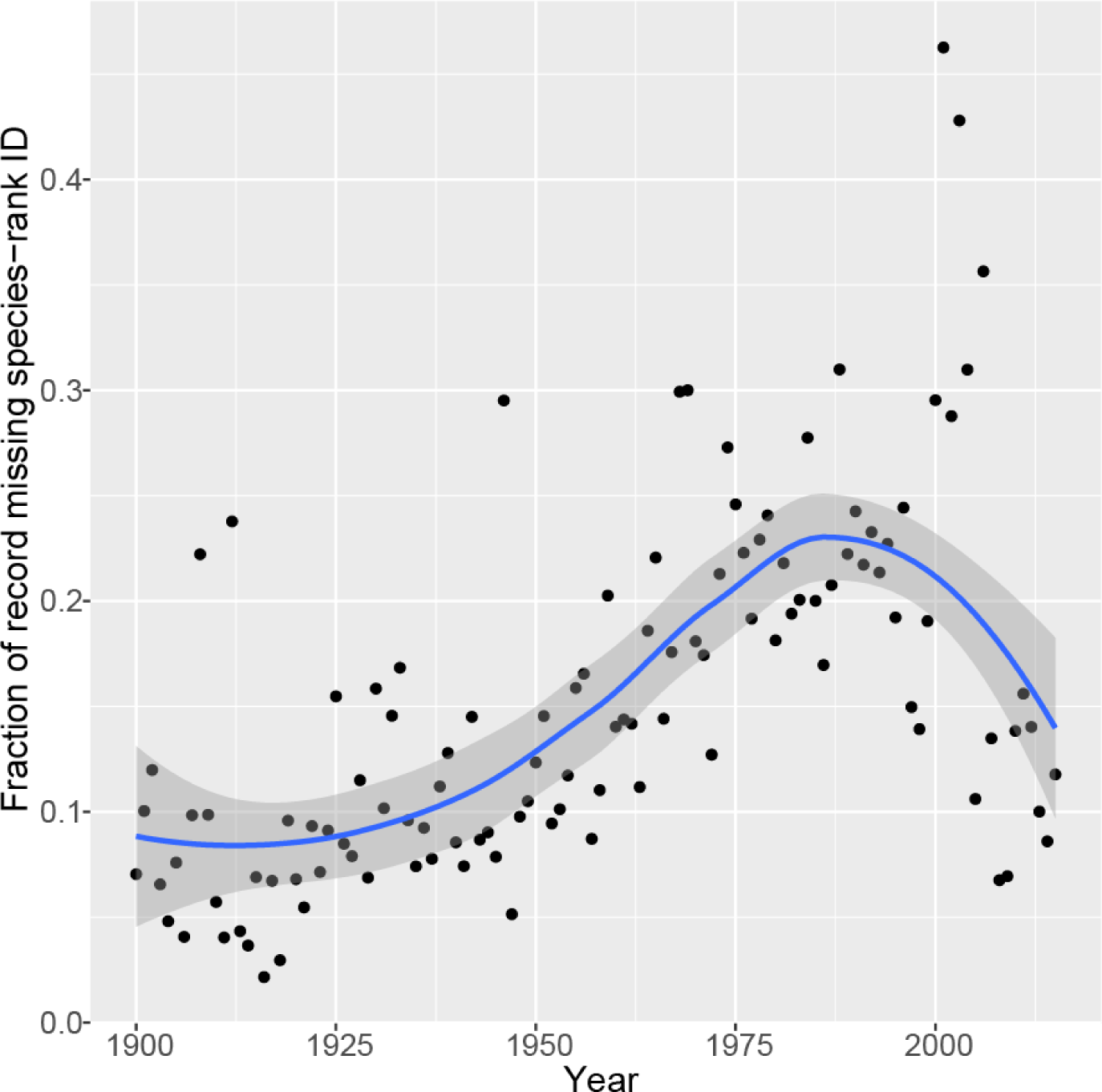
Fraction of the dataset records that lack a species ID. Points show the proportion of records unidentified at the species level in a given year, relative to the total number of records for that year, and the curve shows a loess-smoothed trend line with a smoothing parameter α = 0.75.

**Figure S3:**
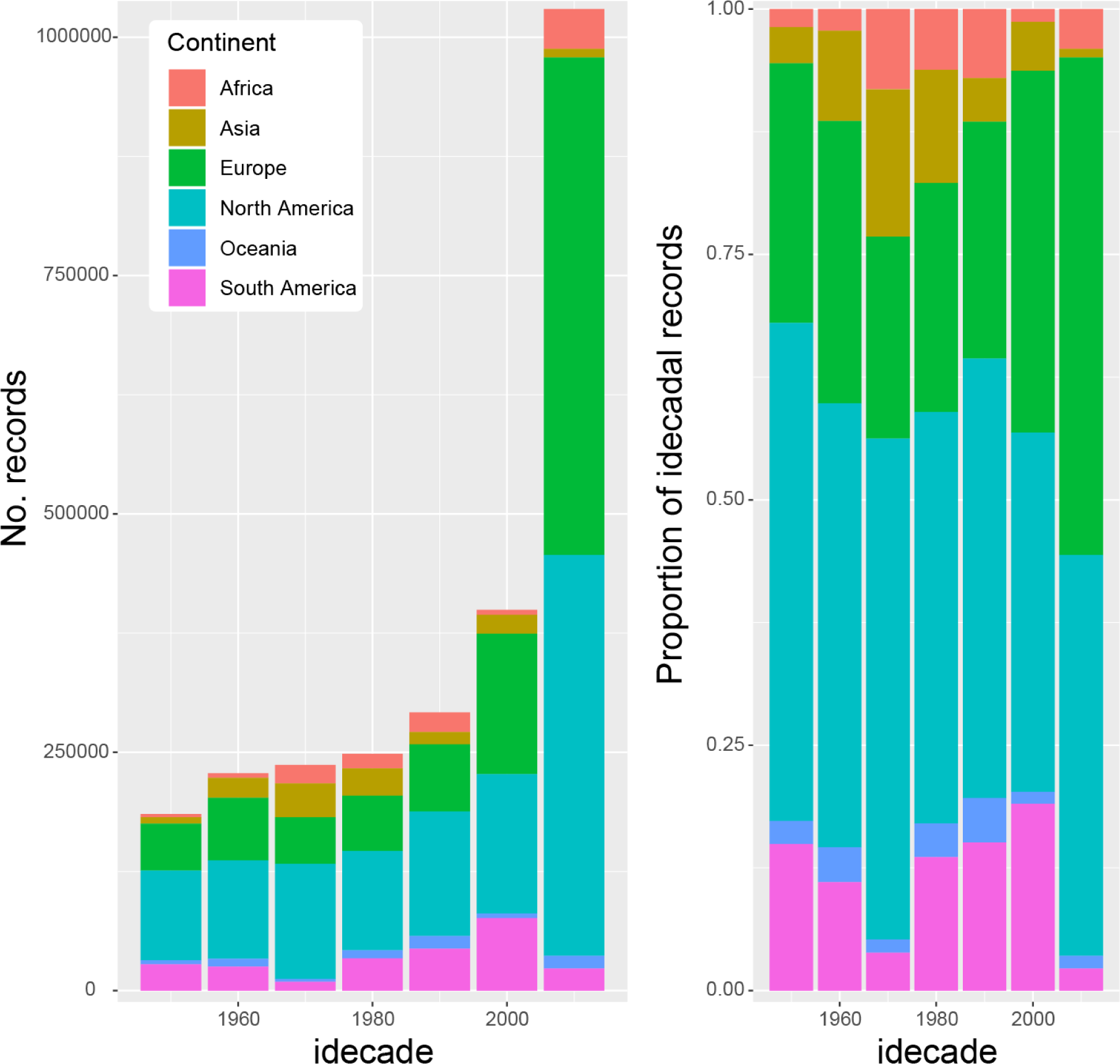
Contribution by idecade of each continent (Antarctica excluded) to the full bee record dataset. (A) Absolute number of GBIF records with a species ID for each continent, grouped by idecade since the 1950’s. (B) Relative contribution of each continent to worldwide idecadal GBIF records with a species ID.

**Figure S4:**
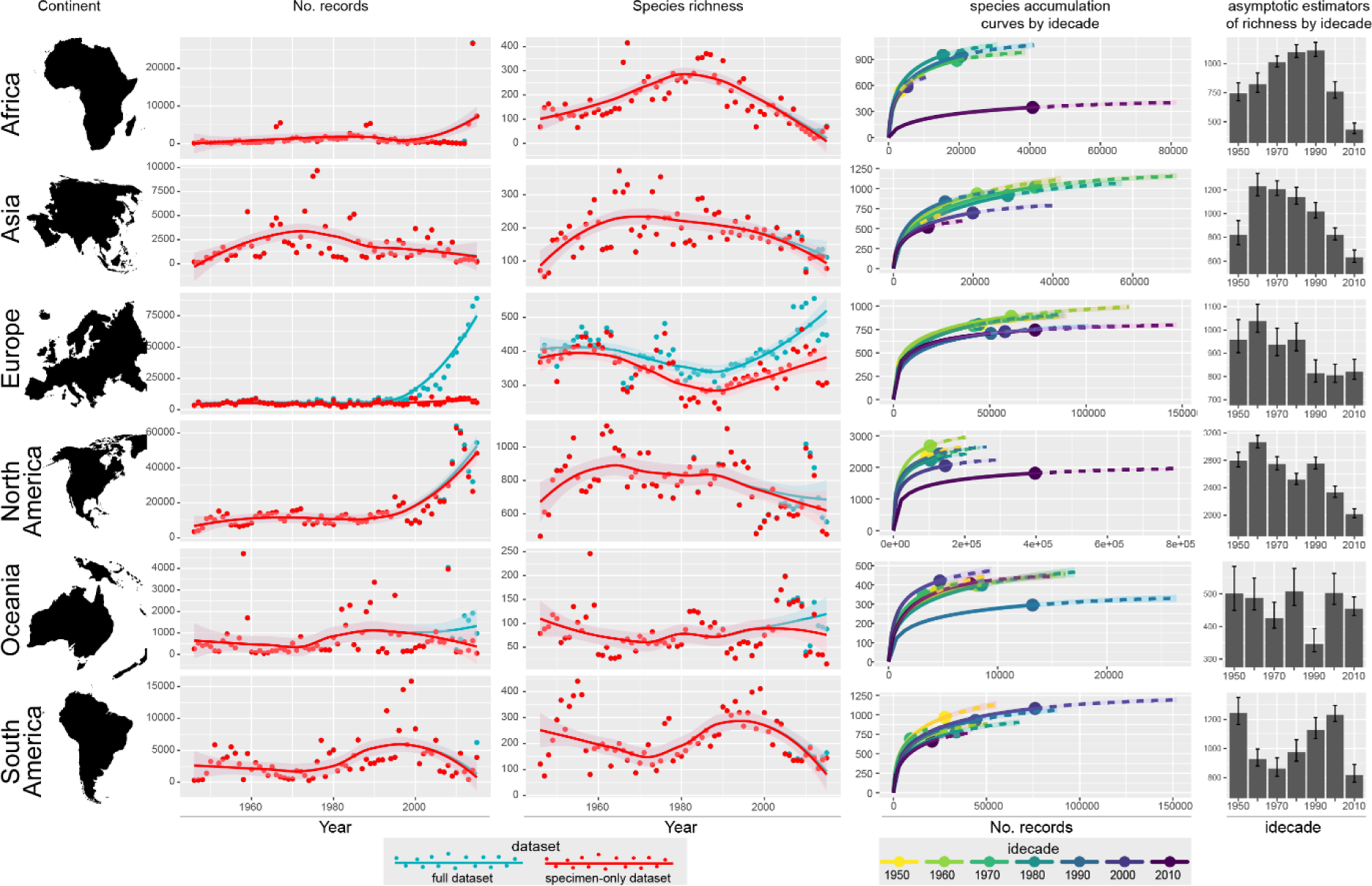
Trends shown in GBIF records for each continent. The left two rows of plots show number of yearly bee records and species in GBIF (blue: full dataset; red: specimens-only dataset); the right two rows show Chao’s interpolation/extrapolation curves based on the specimens-only dataset grouped every ten years (idecades) for the period 1946-2015 and bar plots of the asymptotic estimates of richness by idecade for the same period (error bars mark upper and lower 95% confidence intervals).

**Figure S5:**
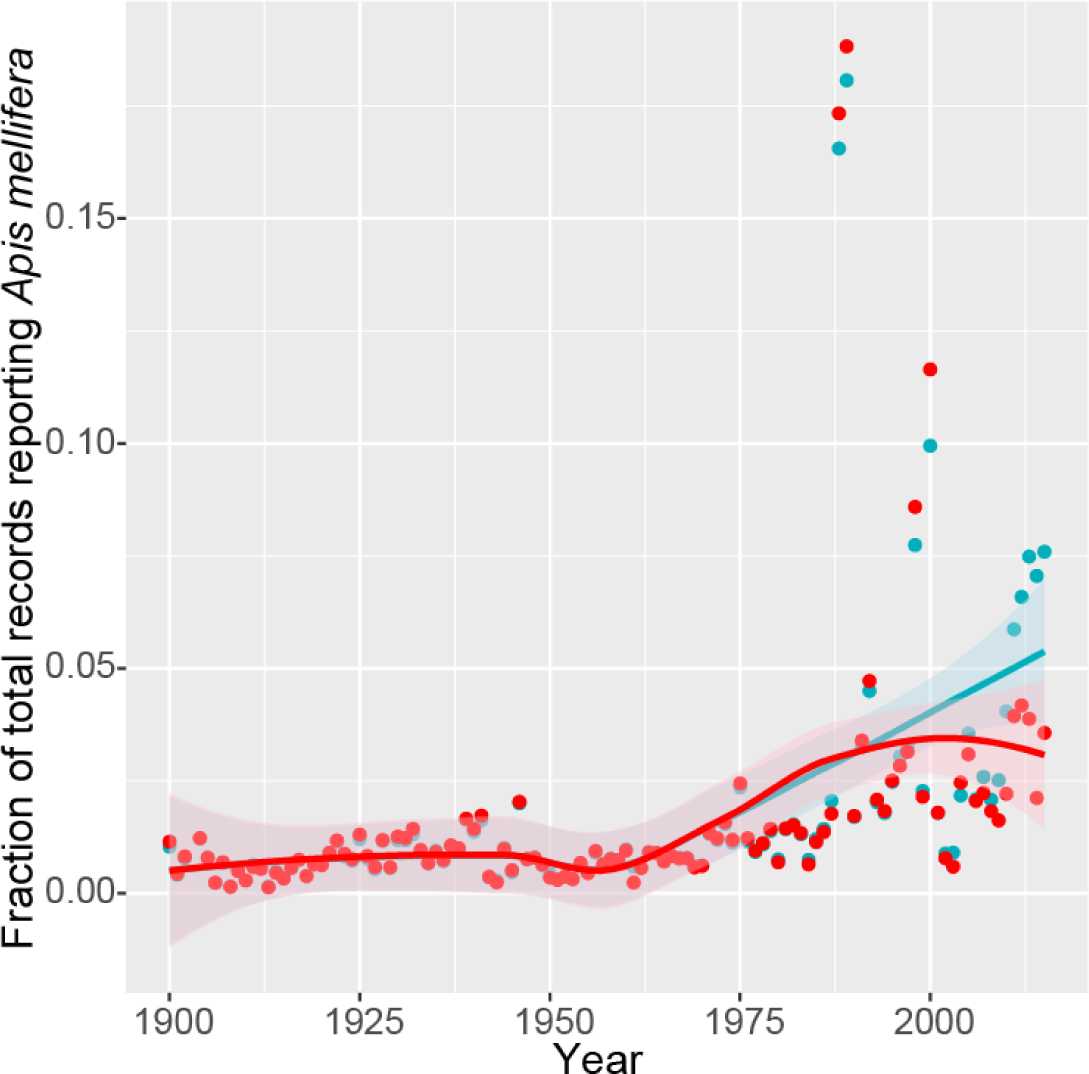
Increase in the fraction of global records of preserved specimens at GBIF represented by the honeybee *Apis mellifera* since the year 1900 (blue: full dataset; red: specimens-only dataset). Points represent yearly proportion of total records belonging to *A. mellifera*; lines show respective loess fit curves with a smoothing parameter α = 0.75.

**Figure S6:**
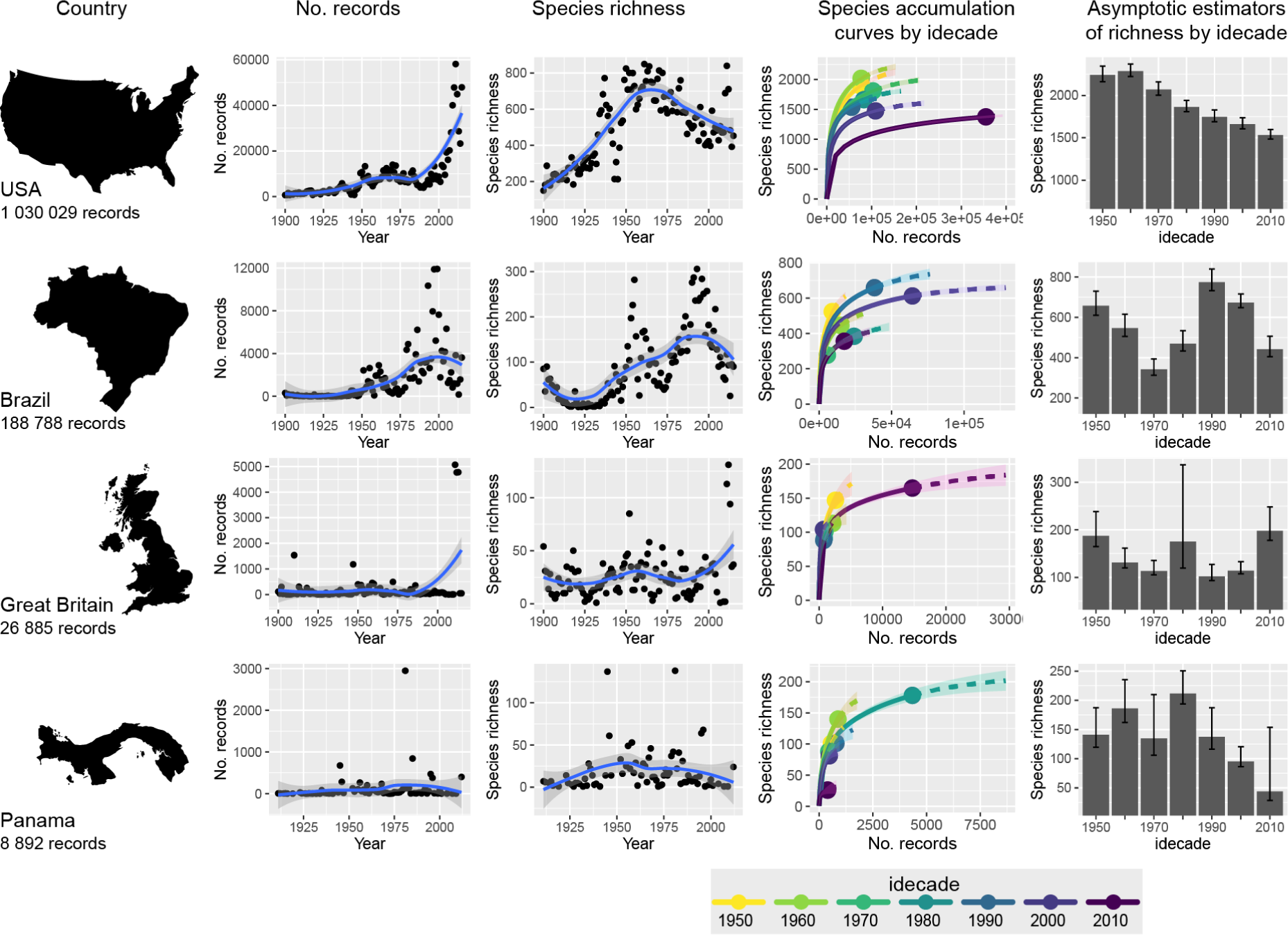
Reliability of trends shown in records of GBIF preserved specimens for specific countries increases with the number of records. The left two rows of plots show number of yearly bee records and species in GBIF for each country – fitted trends are loess curves with a smoothing parameter α = 0.75; the right two rows show Chao’s interpolation/extrapolation curves based on records grouped every ten years (idecades) for the period 1946-2015 and bar plots of the asymptotic estimates of richness by idecade for the same period (error bars mark upper and lower 95% confidence intervals).

